# Variation in mtDNA haplotypes suggests a complex history of reproductive strategy in *Cannabis sativa*

**DOI:** 10.1101/2020.12.28.424591

**Authors:** Ziv Attia, Cloe S. Pogoda, Daniela Vergara, Nolan C. Kane

## Abstract

*Cannabis* is one example in angiosperms that appears to have a recent origin of dioecy and X/Y sex chromosomes. Several evolutionary explanations for this transition have been proposed, with the most parsimonious beginning with a mitochondrial mutation leading to cytoplasmic male sterility (CMS). Our study utilized 73 *Cannabis sativa* whole genome shotgun libraries to reveal eight different mtDNA haplotypes. The most common haplotype contained 60 of the 73 individuals studied and was composed of only dioecious individuals. However, other haplotypes contained a mix of dioecious and monoecious individuals, so haplotype alone does not predict dioecy. From these haplotype groupings we further examined the fully annotated mitochondrial genomes of four hemp individuals and looked for genetic variation affecting reproductive strategy (e.g., monoecious vs. dioecious strategies). Specifically, we searched for markers associated with CMS and for gene rearrangements within these mitochondrial genomes. Our results revealed highly syntenic mitochondrial genomes that contained approximately 60 identifiable sequences for protein coding genes, tRNAs and rRNAs and no obvious rearrangements or chimeric genes. We find no clear evidence that the different reproductive patterns are due to easily identifiable CMS mutations. Our results refute the simplest hypothesis that there was a single recent origin of dioecy in a monoecious ancestor. Instead, the story of the evolution of dioecy is likely much more complex. Further exploration of the nuclear and mitochondrial genomes and their interaction is required to fully understand *Cannabis*’ mating strategies and the possible existence of CMS in this species.

## INTRODUCTION

*Cannabis sativa* is an important annual herb which has been cultivated by humans for millennia. It has extensive amounts of phytochemicals that are used in folk medicine (drug type *Cannabis*) and also contains cellulosic fibers (hemp type *Cannabis*), which are valuable in the textile industry (Russo, 2011; Skoglund et al., 2013, Andre et al., 2016). Presently in the United States, the plant is being developed for more extensive industrial purposes after the approval of the 2018 Agriculture Improvement Act (Congress 2018). Prejudices surrounding this crop species are lessening and research activity continues to increase. Thus, understanding the nature of the *Cannabis* plant as a modern agricultural crop (i.e., hemp) will help to inform the development of it as a valuable plant.

Even though most cultivated *Cannabis* for medical and recreational purposes is dioecious, i.e., male, and female flowers develop on separate plants when grown from seeds, monoecious (i.e., hermaphrodite) populations where male and female flowers are present on the same plant also exist, particularly in the hemp varieties used for industrial purposes. Interestingly, it is unknown what form the wild ancestor took and whether humans have selected for dioecy or monoecy. Most researchers agree that there are no longer any wild populations that can be examined (Small & Cronquist 1976; McPartland 2018). However, modern monoecious varieties have been obtained by selection from naturally occurring variants (Sengbusch et al., 1952; Westergaard 1958; Bocsa 1958). These monecious varieties offer several agronomic / industrial advantages when compared to dioecious cultivars, such as higher crop homogeneity and increased seed yield due to self-fertilization during breeding (Faux et al., 2016, Salentijn et al., 2019). However, monoecy is also associated with some drawbacks, mainly due to inbreeding reducing genetic variation, leading to lower vigor, and slower breeding improvement (Bocsa & Karus 1998). Additionally, when plants produce seeds, they usually devote energy to that process instead of in the production of secondary metabolites such as THCA (delta 9 tetrahydrocannabinolic acid) and CBDA (cannabidiolic acid), making monecious varieties less desirable for medicinal/recreational uses (Lubell & Brand 2018).

*Cannabis sativa* in the haploid state has nine autosomes and one sex chromosome (either X or Y; Mandolino et al., 1999). Like humans, females are typically XX and males are XY. Hermaphrodites also exist, and look cytologically like females, XX (Van Bakel et al., 2011; Razumova et al., 2016). This is not unusual in plants, where sex determination can be influenced by cytoplasmic male sterility (CMS). This means that even though a plant might have the nuclear genome of a hermaphrodite, it is phenotypically female, and does not express male characteristics due to male sterility, perhaps caused by mutations present in the mitochondria (Charlesworth, 2002). CMS is the presence of sterile male reproductive organs (i.e., lack of anthers, or infertile pollen) due to mutations inherited within the organellar genomes, usually mitochondria. The factors that determine sex in plants has evolved numerous times, usually from hermaphroditic origins (Charlesworth, 2002). Many theoretical models of the move from hermaphrodites to monoecious individuals suggest both CMS and other genetic factors play a role (Charlesworth, 2002). These mutations allow for the production of viable female flowers only (e.g., a dioecious plant) and prevent the natural production of pollen (e.g., a monoecious plant). These mutations arise frequently in nature (Hanson, 1991; Balk & Leaver 2001; León et al., 2007) and can be strongly favored if the reduction in pollen production is accompanied by any increase in number of seeds (Duvick, 1990; Vavdiya et al., 2013). Plant breeders use these mutations to control crossing among lines in a variety of crops (Touzet & Meyer 2014; Chen & Liu 2014). CMS is a maternally inherited trait that has been documented in more than 150 higher plant species.

Understanding the mechanisms of CMS, therefore, has broad importance to plant biology and agronomy and may shed light on the evolutionary origins and genetic basis of *Cannabis’* flowering patterns. Two close relatives of *Cannabis, Humulus lupulus* and *H. japonicus* have been studied and it has been shown that sex is controlled by the X to autosome ratio (Divashuk et al., 2014). However, given *Cannabis’* illegal status, research has been inhibited and it is unclear specific effects in the determination of sex and flowering patterns in *Cannabis*. Typical CMS mutations are chimeric mitochondrial open reading frames (ORFs), which are generated by rearrangement and recombination of the mitochondrial genomes (Horn et al. 2014). While most of the identified CMS-related chimeric genes have been linked to ATP synthetase (e.g., atp1, atp4, atp6, atp8 and atp9) or cytochrome C oxidase (e.g., cox1, cox2 and cox3; Hanson & Bentolila 2004), variations in DNA sequence within these genes, as well as in their up/down-stream regions have been shown to be closely linked to CMS in many other crop species (Young & Hanson 1987; Köhler et al. 1991). These candidate genes, together, represent plausible contenders to look for underlying and identifying new CMS mutants.

Here, we compare hemp individuals that represent both monoecious and dioecious reproductive strategies. We hypothesize that if a CMS mutation were a key step in the origins of dioecious *Cannabis*, there would exist mitochondrial polymorphisms and rearrangements associated with the observed monecious and dioecious phenotypes in *Cannabis*. Therefore, our goal was to pursue a detailed examination of these mitochondrial genomes, including annotating protein-coding genes, tRNAs and rRNAs, as well as examining synteny to identify rearrangements.

## MATERIALS AND METHODS

### Whole genome shotgun libraries

We used publicly available Whole Genome Shotgun (WGS) libraries (bioproject PRJNA310948) sequenced by Illumina^™^ Nextera (Lynch et al. 2016, Vergara et al. 2019). These genomes have raw read lengths from 100 to 151bp. Detailed information regarding DNA extraction, sequencing, and library preparation are provided in Lynch et al., (2016) and Vergara et al., (2019). These libraries included 73 *Cannabis sativa* individuals, with some cultivars represented multiple times (Carmagnola X 6, Chocolope X 2, Durban Poison X 2, Afghan Kush X 6, Feral Nebraska X 2, and Kompolti X 2; Table S1). Duplicates were included in our subsequent analyses as a positive control.

### Variant calling

Genomic libraries for 73 *Cannabis sativa* individuals, 67 of them identified in Lynch et al., (2016), were processed to remove adapters and low quality reads by using Trimmomatic v0.39 (Bolger et al., 2014) with the following parameters: Illuminaclip: NexteraPE-PE.fa:2:20:10 Leading:20 Trailing:20 Sliding window:4:15 Minlen:100. The resulting FASTQ files were checked for quality using FASTQC (Andrews, 2010). The quality checked, trimmed sequences were then aligned to the *C. sativa* cs10 assembly (GenBank accession GCA_900626175.2) using the Genome Analysis Toolkit (GATK; Van der Auwera et al., 2013). The resulting variant call file (VCF) table was filtered using vcftools (Danacek et al., 2011) to only include SNPs that specifically aligned to the mitochondrion and had quality scores above 100 (--minQ 100; Table S2). The cs10 assembly (Grassa et al., 2018) was utilized as it is currently the most complete, full annotation publicly available for *Cannabis* and allowed for the entire whole genome libraries to be aligned, which avoided spurious sequence alignment due to similarity between reads for the plastids and nuclear genome.

### Haplotype determination

To determine the major haplotype groups of the 73 *Cannabis* individuals, we used the 1,356 SNPs identified between the 73 *Cannabis* individuals that were present in the filtered VCF table. These SNPs were converted into a FASTA consensus sequence using vcf2phylip.py (Ortiz, 2019; Table S3). The resulting multi-FASTA was analyzed using the R package *pegas* (Paradis, 2010) in R version 3.5.3 to calculate and plot the unique haplotype groups. Each haplotype group was colored based upon reproductive type (Dioecious, Monecious and Unknown). Each of the 73 individuals were assigned to a haplotype group 1-8 (Table S1).

### Genome assembly

In order to carefully determine possible differences between the mitochondrial genomes of monoecious and dioecious *Cannabis* individuals, we focused most of our efforts on two representative monecious and two representative dioecious cultivars. Two of these, Carmagnola (GenBank accession KR059940.1) and Sievers Infinity (GenBank accession KU363807.1) were already assembled / annotated and were obtained from NCBI. The other two, Kompolti (MT361981.1) and Euro Oil (MT557709) were newly assembled and annotated here. In order to assemble Kompolti, and EuroOil, *de novo* assembly of trimmed reads into scaffolds was performed with SPAdes v3.11.1 (Bankevich et al., 2012). Relative position, order and orientation of scaffolds were determined by comparison to available Carmagnola reference genomes. We selected contigs based on read coverage when multiple contigs represented the same genomic region. Contigs were then placed in the correct order and combined by trimming overlapping sequences. Gaps between scaffolds were filled with either raw or trimmed reads that overlapped (e.g., tiling) from the FASTQ files. Once assembled, Zpicture was used to validate the assembly and visualize any potential major differences in structure between the reference (Carmagnola) and each assembled mitochondrial genome (Ovcharenko et al. 2004). Additionally, samtools *tView* was used to confirm the presence of high-quality SNPs (single nucleotide polymorphisms) and/or INDELs (insertions and/or deletions) in the genome. If the mapped reads supported these assembly errors, modifications were made to the FASTA files as needed (Li, 2011).

### Mitochondrial genome annotation

Annotations of genomic features (protein coding sequences, tRNAs and rRNAs) were initiated using GeSeq (Tillich et al., 2017) to find approximate locations of the predicted gene features. In order to identify all possible tRNAs, we utilized tRNAscan-SE 2.0. Additionally, genes not automatically identified using GeSeq were found by using nucleotide and translated protein sequences that were extracted from the reference Carmagnola mitochondrial genome from NCBI. We utilized BLAST (blastn and blastx) to identify regions with homology to these known sequences in our newly assembled FASTA sequences and subsequently annotated any missing features. Annotations were then completed in NCBI’s Sequin 15.50 (Bethesda, MD) and submitted to GenBank for publication.

### Comparative synteny analysis

The comparative positions of genes and reorganization within mitochondrial genomes based on orthologous relationships were plotted using the GUI program MAUVE with default settings (Darling et al., 2004). Comparisons were made between two representative dioecious (Carmagnola and Kompolti) and two monoecious (Sievers Infinity and Euro Oil) individuals.

### Copy number variation

To interrogate the copy number variation (CNV) between two representative monoecious and dioecious individuals, we divided the calculated coverage by depth at every position in the genome. Specifically, we determined coverage by reporting the number of mapped reads (samtools view −c −f) and the depth at every position in the sorted.bam file (samtools depth). The coverage was the calculated as (# mapped reads)*(read length)/(Mitochondrial genome length). Read length was 150 bp for all four haplotypes. CNV was then determined from the formula (coverage)/(depth at every position). These values were then normalized by dividing the CNV at each position by the average of the CNV of the entire genome. Next, we utilized the GenBank annotation files to denote the boundaries of each gene and their associated exons. We report the normalized CNV values for each gene’s exon in Table S4.

## RESULTS

### Variant calling and Haplotype network for 73 Cannabis individuals

Alignment of the 73 *Cannabis sativa* individuals to the cs10 assembly reference genome (as it is currently the most complete publicly available assembly; Grassa et al., 2018) identified a total of 1,356 SNPs (Table S2). Haplotype network prediction using a consensus sequence based on the SNPs (Table S3) produced eight distinct groups. The haplotype groupings are colored based upon sex strategy (Fig. 1; Table S1). The largest group contained 60 of the 73 individuals and showed only the dioecious sex strategy (there were also five individuals that the sex strategy is not yet determined). However, this haplotype grouping suggests that those unknown individuals are likely also dioecious. The second largest group contained six individuals of which two (Dagastani hemp and Sievers Infinity) employed a monoecious sex strategy and four (Finola, Feral Nebraska X 2 and Lebanese) were dioecious. Duplicate cultivars (Carmagnola X 6, Chocolope X 2, Durban Poison X 2, Afghan Kush X 6, Feral Nebraska X 2 and Kompolti X 2) were assigned to the same haplotype groups (group I, I, I, I, II and IV respectively) as positive controls. Given this haplotype map and our interest in determining if the mitochondrial genome affects sex strategy, we focused our analysis on four different hemp individuals. We chose two representative monoecious and two representative dioecious genomes for our subsequent analyses (Figure S1). The first individual was Carmagnola as it was part of the major haplotype group I and is known to be strongly dioecious (Small, 2016). The second dioecious individual selected was Kompolti as it is the most distinct representative of that sex strategy based on our haplotype network. Similarly, we chose Sievers Infinity as a representative monoecious individual, because it is part of the second largest haplotype, group II, and Euro Oil as it is the most distinct monoecious individual based on our haplotype groupings.

**Fig. 1:**
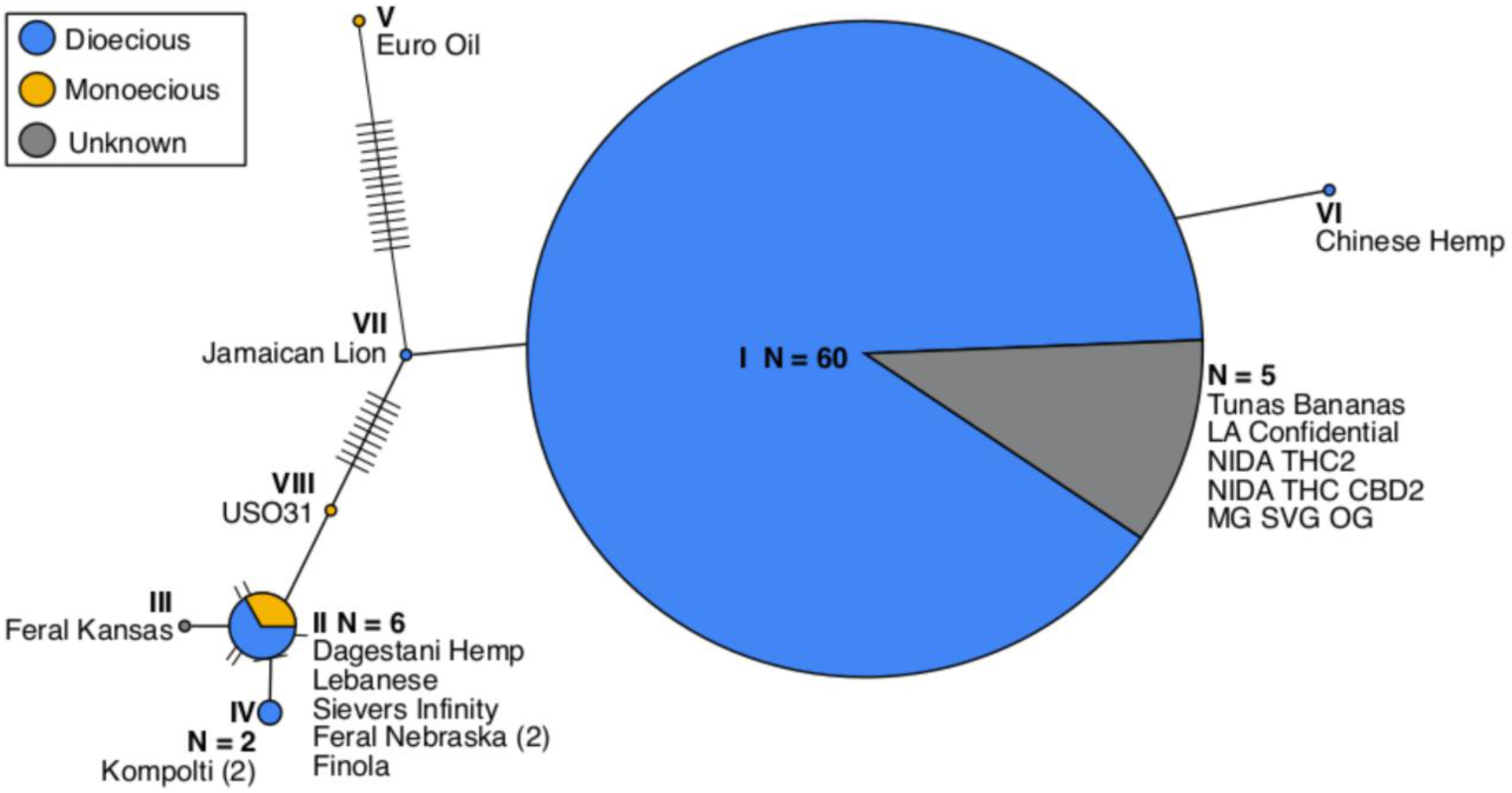
Haplotype network of 73 *Cannabis* individuals (duplicate cultivars are included in the sample size N). The total number of individuals (N), or the individual’s name is given if it is a group of one. Dioecious sex strategy is represented by blue, monecious by yellow and unknown is given in grey.

### Synteny

To begin investigating the possible role of rearrangements of the mitochondrial genome in the CMS phenotype, we compared synteny between our four representative individuals. Synteny is shown in Fig. 2 and is well conserved among the four hemp haplotypes. We specifically examined Carmagnola (Dioecious) vs. Kompolti (Dioecious), Kompolti (Dioecious) vs. Euro Oil (Monoecious), and Euro Oil (Monoecious) vs. Sievers Infinity (Monoecious). All four genomes were aligned to begin with the same nucleotide sequence to ensure appropriate alignment of the genomes. It is clear that there are no rearrangements between any of individuals and specifically no major or minor difference between the dioecious and monoecious groups was observed (Fig. S2).

**Fig 2.**
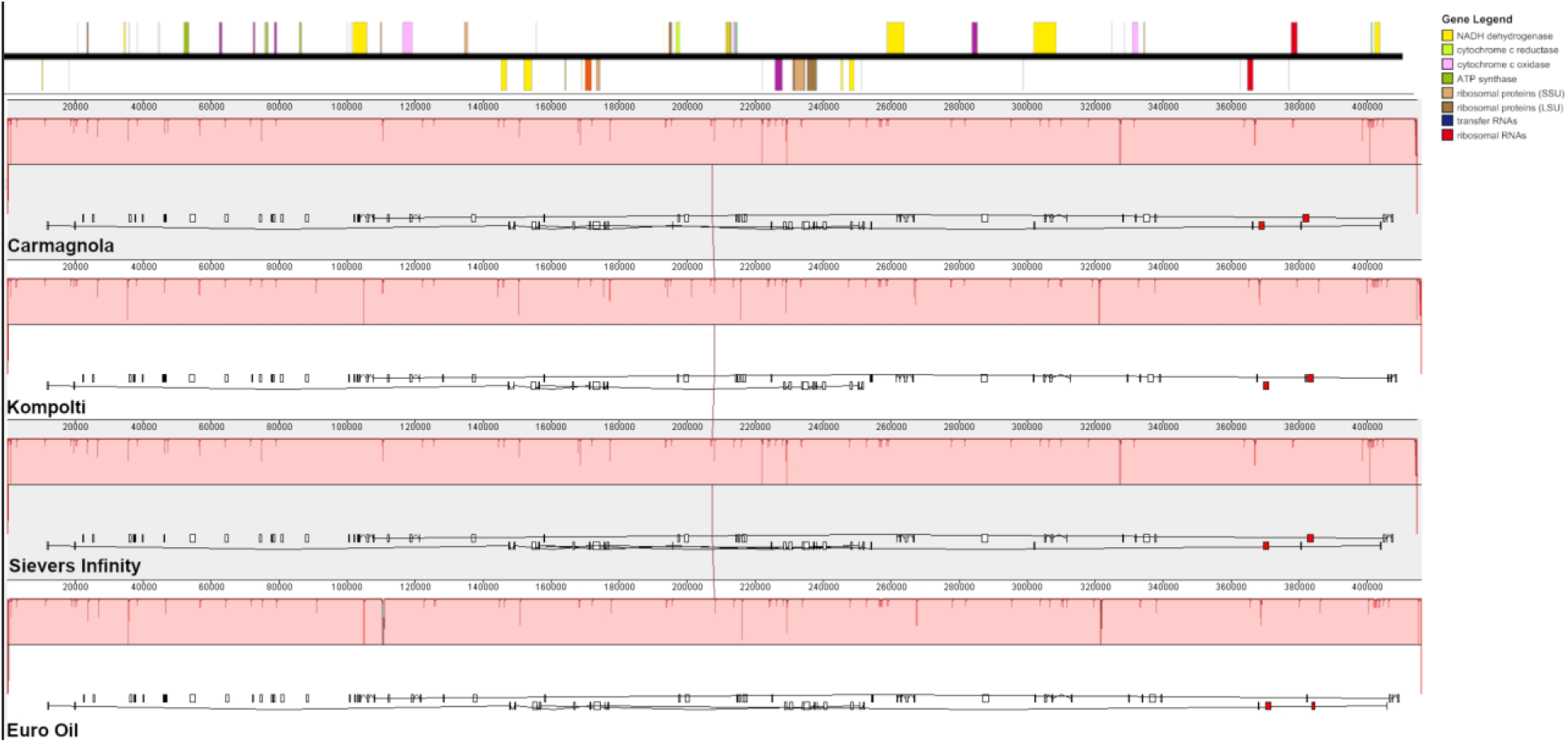
Synteny plot of four hemp haplotypes: Carmagnola (KR_059940), Kompolti (MT361981.1), Euro Oil (MT557709) and Sievers infinity (KU363807.1). Genetic content for the reference (Carmagnola) is shown at the top of the figure and the figure legend (top right) shows what each colored object represents. All haplotypes were of similar size (400 kb) and they are aligned to each other.

### Mitochondrial genomic content

In order to shed light on the differences in flowering pattern strategies of the *Cannabis* mitochondrial genomes used here, we compared four different hemp mitochondrial genomes: two representative monoecious and two representative dioecious individuals. The mitochondrial gene content was highly similar between the four mitochondrial genomes we analyzed. We identified approximately 60 genes in each mitochondrial genome including protein coding genes, tRNAs and rRNAs. Forty-two protein coding genes previously associated with the CMS phenotype were present in all these genomes and are closely compared in Table 1, with only minor observable differences in some cases, such as different numbers of exons or strand orientation of a given gene. These differences however, could be due to slight variability in annotation rather than underlying genetic differences because genome-wide, nucleotide similarity for the assembled mitochondrial genomes (i.e., FASTA sequences) was 100% for Carmagnola vs. Sievers Infinity, 99.94% for Carmagnola vs. Kompolti, and 99.96% for Carmagnola vs. Euro Oil. The differences between dioecious (Carmagnola) and monecious (Sievers Infinity and Kompolti) and whether they occurred in a coding region or not, are presented in Table 2. Out of 25 potential SNPs and INDELs that were different between the Carmagnola representative dioecious individual and both our monoecious individuals (Euro Oil and Sievers Infinity), there were only five that occurred near an annotated gene, with only one (an insertion of AG at position 26,7314 bp) occurring in an exon of nad7 in Carmagnola. However, alignment of this protein did not reveal any amino acid changes, as this insertion occurs outside the coding regions in both the dioecious individuals. GC percentage was identical between the four haplotypes and was found to be 45.6 %.

**Table 1:**
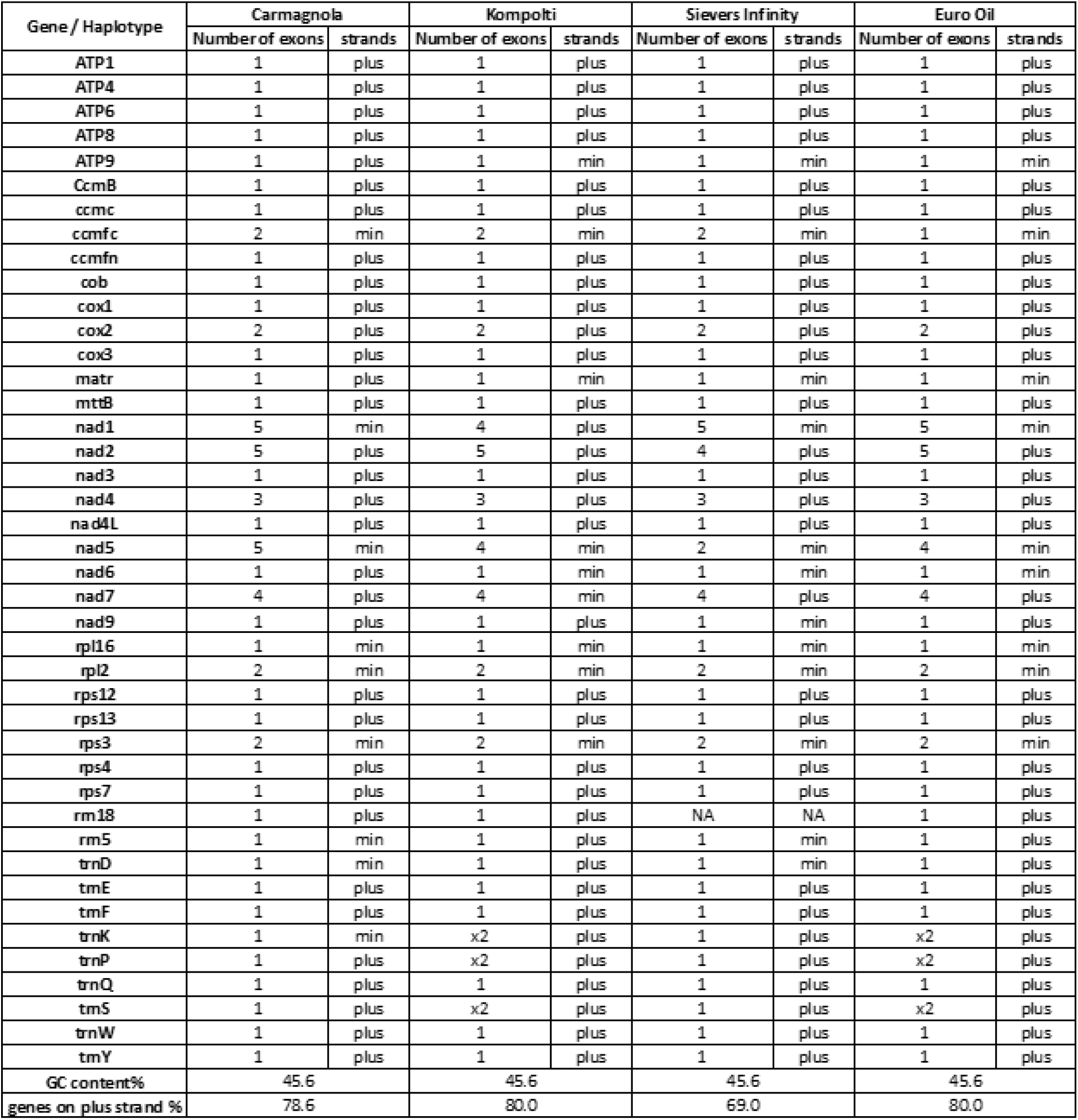
Gene presence / absence for the two representative dioecious individuals (Carmagnola and Kompolti) and two representative monecious individuals (Sievers Infinity and Euro Oil). Number of exons for each gene, tRNA, and rRNA are given as well as if it was on the plus or minus strand. Percentage of coding sequence on the plus strand and GC content is given for each genome.

| Gene / Haplotype | Carmagnola |  | Kompolti |  | Sievers Infinity |  | Euro Oil |  |
| --- | --- | --- | --- | --- | --- | --- | --- | --- |
|  | Number of exons | strands | Number of exons | strands | Number of exons | strands | Number of exons | strands |
| ATP1 | 1 | plus | 1 | plus | 1 | plus | 1 | plus |
| ATP4 | 1 | plus | 1 | plus | 1 | plus | 1 | plus |
| ATP6 | 1 | plus | 1 | plus | 1 | plus | 1 | plus |
| ATP8 | 1 | plus | 1 | plus | 1 | plus | 1 | plus |
| ATP9 | 1 | plus | 1 | min | 1 | min | 1 | min |
| CcmB | 1 | plus | 1 | plus | 1 | plus | 1 | plus |
| ccmc | 1 | plus | 1 | plus | 1 | plus | 1 | plus |
| ccmfc | 2 | min | 2 | min | 2 | min | 1 | min |
| ccmfn | 1 | plus | 1 | plus | 1 | plus | 1 | plus |
| cob | 1 | plus | 1 | plus | 1 | plus | 1 | plus |
| cox1 | 1 | plus | 1 | plus | 1 | plus | 1 | plus |
| cox2 | 2 | plus | 2 | plus | 2 | plus | 2 | plus |
| cox3 | 1 | plus | 1 | plus | 1 | plus | 1 | plus |
| matr | 1 | plus | 1 | min | 1 | min | 1 | min |
| mtt8 | 1 | plus | 1 | plus | 1 | plus | 1 | plus |
| nad1 | 5 | min | 4 | plus | 5 | min | 5 | min |
| nad2 | 5 | plus | 5 | plus | 4 | plus | 5 | plus |
| nad3 | 1 | plus | 1 | plus | 1 | plus | 1 | plus |
| nad4 | 3 | plus | 3 | plus | 3 | plus | 3 | plus |
| nad4L | 1 | plus | 1 | plus | 1 | plus | 1 | plus |
| nad5 | 5 | min | 4 | min | 2 | min | 4 | min |
| nad6 | 1 | plus | 1 | min | 1 | min | 1 | min |
| nad7 | 4 | plus | 4 | min | 4 | plus | 4 | plus |
| nad9 | 1 | plus | 1 | plus | 1 | min | 1 | plus |
| rpl16 | 1 | min | 1 | min | 1 | min | 1 | min |
| rpl2 | 2 | min | 2 | min | 2 | min | 2 | min |
| rps12 | 1 | plus | 1 | plus | 1 | plus | 1 | plus |
| rps13 | 1 | plus | 1 | plus | 1 | plus | 1 | plus |
| rps3 | 2 | min | 2 | min | 2 | min | 2 | min |
| rps4 | 1 | plus | 1 | plus | 1 | plus | 1 | plus |
| rps7 | 1 | plus | 1 | plus | 1 | plus | 1 | plus |
| rm18 | 1 | plus | 1 | plus | NA | NA | 1 | plus |
| rm5 | 1 | min | 1 | plus | 1 | min | 1 | plus |
| trnD | 1 | min | 1 | plus | 1 | min | 1 | plus |
| tmE | 1 | plus | 1 | plus | 1 | plus | 1 | plus |
| tmF | 1 | plus | 1 | plus | 1 | plus | 1 | plus |
| trnK | 1 | min | x2 | plus | 1 | plus | x2 | plus |
| trnP | 1 | plus | x2 | plus | 1 | plus | x2 | plus |
| trnQ | 1 | plus | 1 | plus | 1 | plus | 1 | plus |
| tmS | 1 | plus | x2 | plus | 1 | plus | x2 | plus |
| trnW | 1 | plus | 1 | plus | 1 | plus | 1 | plus |
| tmY | 1 | plus | 1 | plus | 1 | plus | 1 | plus |
| GC content% | 45.6 |  | 45.6 |  | 45.6 |  | 45.6 |  |
| genes on plus strand % | 78.6 |  | 80.0 |  | 69.0 |  | 80.0 |  |

**Table 2:** SNP and INDEL differences between the reference Dioecious individual (Carmagnola) and the alternate Monecious individuals (Euro Oil and Sievers Infinity). Position is given in bp for the reference Carmagnola annotation. The gene feature corresponds to the reference Carmagnola annotation.

| Position (bp) in Carmagnola | Reference (Dioecious) | Alternate (Monoecious) | Gene Feature in Carmagnola |
| --- | --- | --- | --- |
| 3,208 | C | T | Non-Coding |
| 56,571 | C | A | Non-Coding |
| 74,900 | CT | C | Non-Coding |
| 122,196 | C | A | Non-Coding |
| 144,424 | A | G | Non-Coding |
| 150,582 | G | T | Non-Coding |
| 167,988 | C | A | Intron trans-spliced nad2 |
| 213,604 | A | T | 5' Leader sequence nad3 |
| 247,699 | C | G | 5' Leader sequence nad6 |
| 256,390 | T | G | Non-Coding |
| 257,100 | C | G | Non-Coding |
| 267,314 | CAG | CAGAG | Exon nad7 |
| 277,630 | T | G | Non-Coding |
| 295,239 | A | T | Non-Coding |
| 306,408 | T | C | Intron nad4 |
| 318,064 | C | A | Non-Coding |
| 331,459 | A | C | Non-Coding |
| 363,735 | T | G | Non-Coding |
| 378,146 | A | C | Non-Coding |
| 384,437 | AGGG | AGGGG | Non-Coding |
| 400,445 | A | C | Non-Coding |
| 401,584 | C | G | Non-Coding |
| 404,499 | G | T | Non-Coding |
| 413,301 | ATC | ATCGTTC | Non-Coding |
| 413,302 | TC | TCGTCC | Non-Coding |

### Copy number variation

Copy number variation (CNV) was similar across the monoecious and dioecious hemp individuals and 42 genes analyzed (Fig. 3). The average CNV was approximately ~1 and is not likely contributing to sex determination or the CMS phenotype. The only outlier was the tRNA trnP (Proline), which has much higher copy number only in the Carmagnola mitochondrial genome.

**Fig 3.**
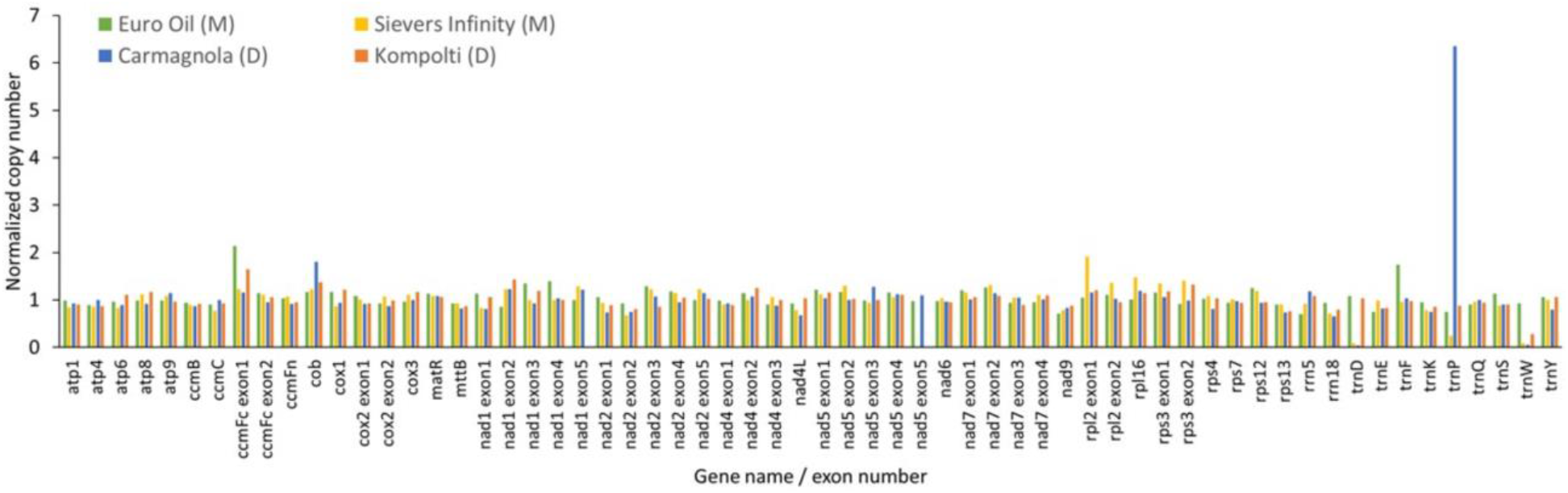
Copy number variation (CNV) for Carmagnola, Sievers infinity, Kompolti and Euro Oil hemp haplotypes. Values were normalized by dividing by the average CNV of each individual mitochondrial genome.

## DISCUSSION

This study takes a detailed look at different *Cannabis* haplotypes to elucidate differences in the mitochondrial genomes for these various cultivars. Using 73 *Cannabis sativa* individuals aligned to the cs10 (Grassa et al. 2018) reference genome, we identified eight distinct haplotypes, with the majority of the dioecious individuals (60) falling within haplotype group I (Fig. 1). We chose to closely examine four separate individuals, two (Kompolti and Euro Oil) of which were newly assembled and annotated here. The first of the four we examined, Carmagnola, was part of the main haplotype group I and was dioecious. The second, Sievers Infinity, fell within haplotype group II and was monoecious. Additionally, to increase our success in revealing differences in sex determination we chose the most distinct dioecious (Kompolti) and monecious (Euro Oil) individuals as well. All individuals examined are hemp varieties of *Cannabis sativa* and are cultivated for industrial purposes. Carmagnola is strongly dioecious, existing always as separate male and female plants. Kompolti is also dioecious but can express flowers of either sex on the same plant, with the use of chemicals (e.g., Ag^+^ ions and gibberellins; Ram & Jaiswal 1972; Atsmon & Tabbak 1979; Ram & Sett 1982). In contrast, Sievers infinity and Euro oil are monoecious (Table S1). Our analyses revealed that the mitochondrial genomes of all four individuals were remarkably similar and did not demonstrate any obvious signs or patterns of cytoplasmic male sterility (CMS). The genetic content, SNPs/INDELs, CNV and synteny were all highly conserved. This observation is remarkably similar to previous research that looked for differences in the chloroplast genomes of *Cannabis* and found that they too were highly similar and conserved (Vergara et al., 2015; Roman et al., 2019).

There are two types of sterility possible in plants: mitochondrial or nuclear encoded. Mitochondrial encoded sequences conferring cytoplasmic male sterility is commonly observed in plant species and is the result of genomic conflict between the mitochondrial and nuclear genomes (Horn et al., 2014; Touzet & Meyer 2014). In contrast, nuclear, or genic, male sterility has been associated with 20 nuclear associated gene mutations (Neuffer et al., 1997) and it is also common in flowering plants (Gabay-Laughnan & Laughnan 1994). Given our haplotype group results, we suggest that dioecy does not have a single, recent origin, since similar haplotypes are found in dioecious individuals. Additionally, a single, ancient origin of monoecy from a dioecious ancestor is also not supported. If dioecy were ancestral as suggested by previous research (Kovalchuck et al., 2020) we would expect more diversity in those dioecious individuals and not the single large haplotype observed. Group I of our haplotype network contains 60 individuals and suggests that instead there was a recent selective sweep specific to this mitochondrial locus, but it is not the only cytotype associated with dioecy. The lack of observed diversity in both the mitochondrial and chloroplast genomes are in direct contrast to the nuclear genome diversity in these individuals (Lynch et al. 2016). Thus, we can refute both a recent origin of dioecy as well as a single ancient origin of monoecy. This suggests that instead of a simple CMS mutation controlling reproductive strategy there has been a fairly complex evolution of dioecy vs. monoecy in *Cannabis*, perhaps involving several distinct mutations and likely involving the nuclear genome.

## CONCLUSIONS

As the nature of *Cannabis* continues to evolve towards being an accepted agricultural crop, more information about agronomic traits is required. This crucial information will provide valuable tools for breeding hemp. Currently, most *Cannabis* for medical and recreational purposes propagated vegetatively, and stocks of homogenous seed in the US do not exist. *Cannabis* lacks the genetic and genomic tools available for most important agricultural crops due to its illegal status (Vergara et al., 2016), and the absence of basic resources such as public isogenic germplasm collections hinders the improvement and development of cultivars. Crop improvement specialists require creative and collaborative solutions to overcome these issues due to years of scientific neglect. Therefore, in order to make *Cannabis* more appealing and widely available to commercial farmers, “true breeding” (e.g., F1 hybrids) approaches are necessary. In addition, focused crossing will allow the development of bi-parental populations and introgression lines that facilitate sophisticated genomic approaches, such as genome wide association studies (GWAS). These methods are well established in similar and utilized in other agricultural crops like sunflower, corn, and tomato. (Vear et al., 2016; Bauchet et al., 2017; Darrah et al., 2019). This type of research has important implications for the medical, recreational, and industrial industries that rely on *Cannabis*. Fully understanding the nature of the sterility phenotype will be a huge step forward in the process of making *Cannabis* into a profitable, accessible, and modern crop.

## AUTHOR CONTRIBUTION

Z.A. and C.S.P assembled the two new genomes, wrote the manuscript and created figures. D.V. and N.C.K conceived and consulted on the project. All authors contributed to manuscript preparation.

## DATA ACCESSIBILITY

All mitochondrial genomes are publicly available from NCBI: Carmagnola (KR_059940), Sievers Infinity (KU363807.1), Kompolti (MT361980.1) and Euro Oil (MT557709-temporary accession number, awaiting final publication).

## ACKNOWLEDGMENTS

The authors thank all companies and people who provided DNA samples or sequence information.

