## Supplementary material for "Variation in mtDNA haplotypes suggests a complex history of reproductive strategy in *Cannabis sativa*": sup data

### **Supplementary Information**

**Table S1**: Cultivar name, reproductive type (D for Dioecious, H for Hermaphrodite (monoecious)), and haplotype group number for the 73 individual *Cannabis sativa* samples used here. Duplicate cultivar names are provided. Haplotype group number is 1-8 and corresponds to figure 1 in the main text.

**Table S2**: Variant call format table (SNPs & INDELs) for the 73 *Cannabis sativa* cultivars aligned to the reference cs10 (VCF was created using GATK).

**Table S3**: FASTA consensus sequence for each of the 73 *Cannabis sativa* cultivars aligned to the reference cs10 (Grassa et al. 2018). 1,356 SNPs are included for each individual.

**Table S4**: The normalized CNV values for each gene’s exon for the four hemp haplotypes.

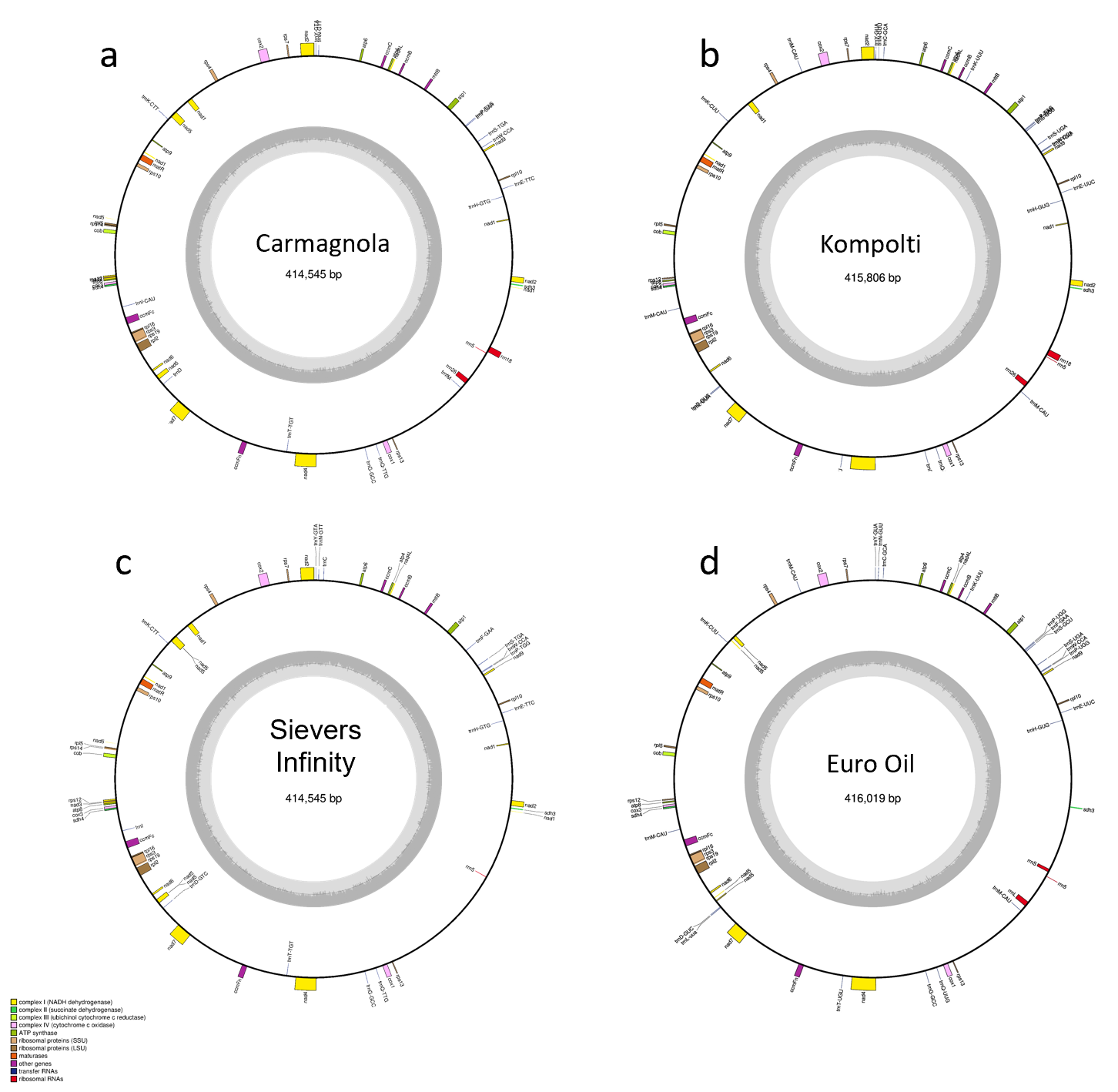

**Fig S1.** Genomic content for the Carmagnola (KR_059940) and Sievers Infinity (KU363807.1) as well as the two newly sequenced, assembled and annotated *Cannabis sativa* hemp Kompolti (MT361980.1) and Euro Oil (MT557709) mitochondrial genomes. Figures were produced using the OGDraw function from GeSeq (Tillich et al., 2017). Genomic content is given in the figure legends.

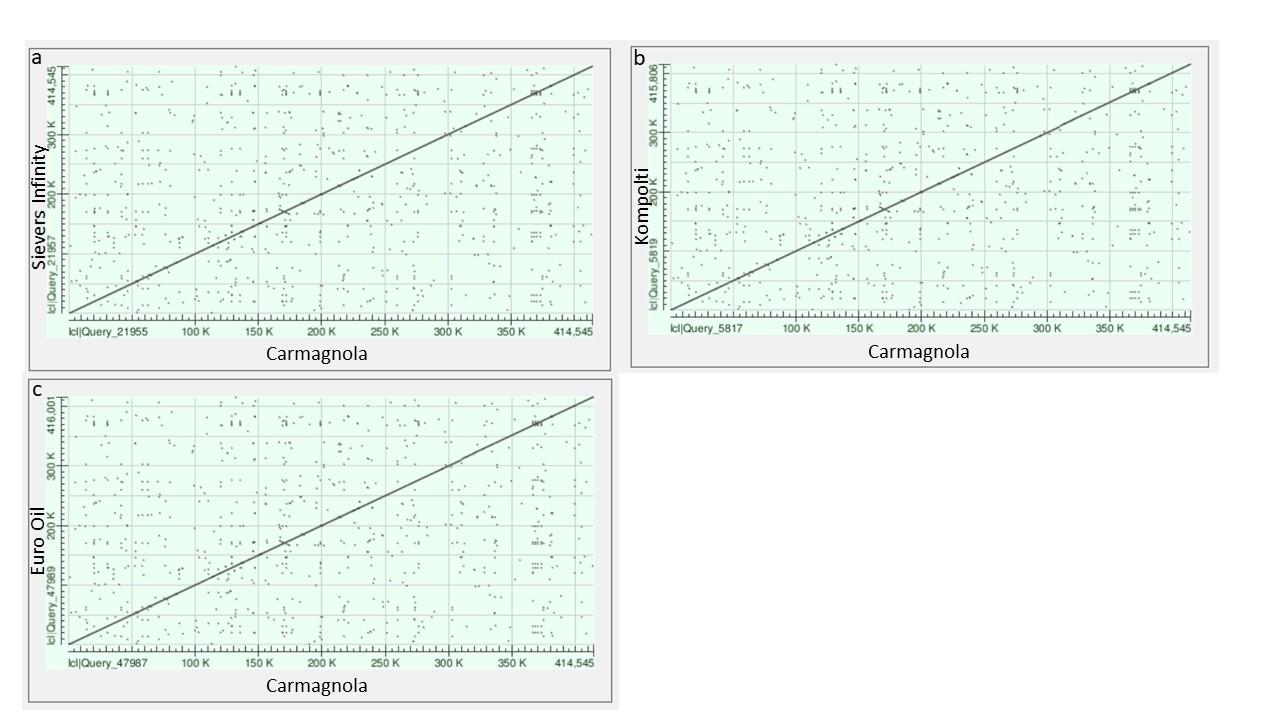

**Fig S2.** Dot plot synteny comparisons performed via NCBI’s blastn. FASTA sequences for Carmagnola (KR_059940) was compared against **(a)** Sievers Infinity (KU363807.1), **(b)** Kompolti (MT361980.1) and **(c)** Euro Oil (MT557709). Figures were produced via NCBI’s blastn web interface.
